# Multimodal Cross-Attentive Graph-Based Framework for Predicting *In Vivo* Endocrine Disruptors

**DOI:** 10.1101/2025.09.16.676584

**Authors:** Eder Soares de Almeida Santos, Gustavo Felizardo Santos Sandes, Artur Christian Garcia da Silva, Holli-Joi Martin, Eugene N. Muratov, Braga Rodolpho de Campos, Bruno Junior Neves

## Abstract

Endocrine hazard assessment needs models that are accurate and mechanistically transparent. We present a multimodal cross-attentive graph framework that fuses molecular graphs with adverse-outcome-pathway (AOP)–anchored assay signals to predict organism-level outcomes in the OECD Hershberger and uterotrophic assays. In Tier-1, multitask GNNs learn ER/AR molecular-initiating and key events across 46 ToxCast/Tox21 assays. In Tier-2, a cross-attentive multimodal GNN integrates Tier-1 pathway signals with molecular graphs, achieving AUROC□=□0.90 (Hershberger) and 0.96 (uterotrophic). External validation on literature compounds showed 84% concordance (Hershberger 15/18; uterotrophic 22/26). Bidirectional cross-attention links molecular substructures to pathway assays and vice-versa, while counterfactual perturbations rank assays and structural motifs most responsible for each decision. The framework couples high accuracy with assay-traceable explanations, supporting targeted testing within the Integrated Approaches.

## 1. INTRODUCTION

Endocrine-disrupting chemicals (EDCs) comprise a group of exogenous substances that interfere with hormone signaling pathways, particularly through estrogen (ER) and androgen (AR) receptors mechanisms,^1,2^ leading to well-documented adverse outcomes ranging from reproductive disorders to hormone-dependent cancers.^3–7^ International evaluations and economic analyses estimate that such exposures contribute to widespread disease at the population level and generate annual costs amounting to hundreds of billions of euros in the European Union.^8^

Assessment of chemical hazards has historically relied on *in vivo* assays established under the Organization for Economic Co-operation and Development (OECD) test guidelines to capture AR- and ER-mediated mechanisms. The Hershberger assay (OECD test no. 441), conducted in castrated male rats supplemented with testosterone propionate, provides direct evidence of AR agonism or antagonism through changes in male accessory sex organ weights. Similarly, the uterotrophic assay (OECD test no. 440), performed in immature or ovariectomized female rodents, monitors ER modulation by measuring uterine weight and histological changes. While these assays serve as regulatory benchmarks for identifying endocrine activity, they have limited throughput, high costs, and ethical concerns associated with the use of animals.^9^

In addition, high-throughput screening strategies have been established for assessing EDCs. The U.S. Environmental Protection Agency’s (EPA) ToxCast program and the interagency Tox21 collaboration have generated extensive *in vitro* bioactivity data across ∼10,000 chemicals,^10–12^ including dedicated assay batteries for ER and AR pathways.^13,14^ These resources provide unprecedented mechanistic insight for regulatory prioritization but remain constrained by limited metabolic competence and the challenge of translating pathway perturbations into complex *in vivo* outcomes.^15,16^ Building on these efforts, the U.S. EPA implemented the Endocrine Disruptor Screening Program (EDSP) to determine potential endocrine effects in humans and wildlife. The program was structured in two tiers: Tier-1 was designed to identify potential EDCs through a battery of *in vitro* and *in vivo* assays (including the OECD Hershberger and uterotrophic tests); and Tier-2, intended to characterize dose-response relationships and adverse outcomes in more comprehensive animal studies.^17,18^ Despite its ambitious scope, the EDSP has progressed slowly, with Tier-1 testing completed for only a limited number of chemicals.

Considering these limitations, computational toxicology has emerged as a promising approach to enhance the throughput of chemical assessments.^19^ Within computational toxicology, machine learning approaches leverage large-scale biological, chemical, and mechanistic data to uncover non-linear patterns underlying endocrine disruption,^20–27^ thereby enabling regulatory prioritization while reducing reliance on animal testing. These approaches have been increasingly adopted into Integrated Approaches to Testing and Assessment (IATA) frameworks as complementary tools to conventional assays, offering a scalable strategy to screen the thousands of chemicals that remain untested for endocrine activity.^28,29^

In recent years, graph neural networks (GNNs)^30,31^ have expanded the landscape of predictive toxicology. These models rely on molecular graphs, where nodes represent atoms, edges represent bonds, and attributes (such as atom type, chirality, and bond order) are encoded as features. GNNs offer adaptable architectures for modeling complex molecular interactions and endocrine-related responses. However, current applications in computational toxicology rely primarily on molecular structure information, with underutilization of biological pathway data. Despite promising advances, these studies remain limited in number and scope, focusing mainly on *in vitro* AR- and ER-mediated effects.^32–34^ Extending these efforts toward more comprehensive predictions, particularly those aligned with organism-level outcomes, requires integrative modeling strategies that combine molecular graph representations with adverse outcome pathways and assay-derived information. Such approaches enable the capture of receptor-level interactions while also accounting for downstream biological responses.

In this study, we introduce a multimodal cross-attentive graph framework that uniquely integrates chemical structure information with adverse outcome pathway (AOP)-based predictions to model *in vivo* endocrine disruption. The approach was developed in two tiers: first, multitask GNNs were trained on 46 *in vitro* assays from the ToxCast/Tox21 program to capture AR- and ER-mediated molecular initiating events (MIEs) and key events (KEs); second, the resulting predictions and logits—rather than the raw *in vitro* labels—were embedded within a pathway graph and integrated with molecular graphs through cross-attentive multimodal GNNs to predict outcomes of the OECD Hershberger (AUROC = 0.90) and uterotrophic (AUROC = 0.96) assays.

Beyond predictive performance, the multimodal framework provides explainability through bidirectional cross-attention, which links molecular substructures to pathway assays and, conversely, pathway nodes to their most influential molecular features. This dual mapping elucidates how structural motifs propagate into AR- and ER-mediated perturbations. As an additional interpretability strategy, counterfactual perturbations were employed to generate alternative scenarios, enabling the evaluation of how molecular modifications or pathway disruptions may alter downstream outcomes.

## 2. RESULTS AND DISCUSSION

### 2.1. Tier-1

Tier-1 modeling was designed to predict *in vitro* responses across AR- and ER-mediated assays, establishing a mechanistic basis for downstream *in vivo* predictions. Initially, a dataset of 7,957 chemicals tested in 46 *in vitro* assays from the U.S. EPA ToxCast/Tox21 program (invitroDB v4.1) was compiled within AR- and ER-mediated AOPs (Figure 1). These assays capture a structured sequence of biological events, linking MIEs such as receptor binding to successive KEs (e.g., transcriptional regulation, gene expression, and cell proliferation) that ultimately culminate in an adverse outcome (AO) at the organismal level.

**Figure 1.**
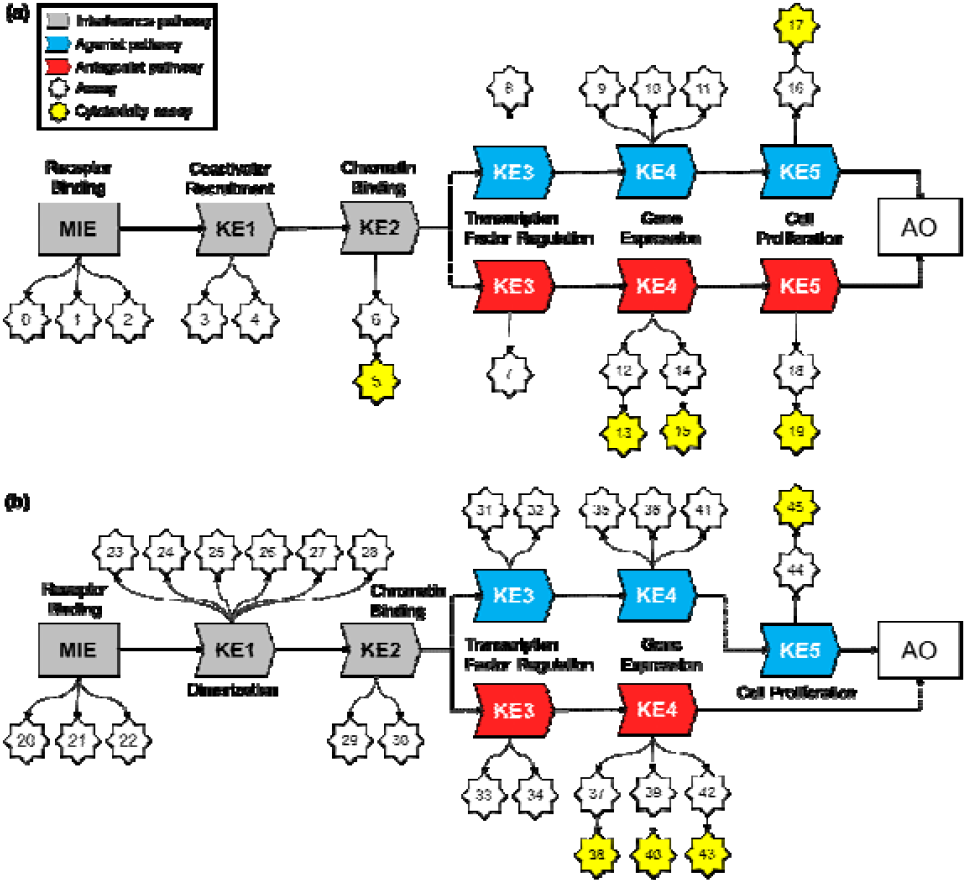
Graphical representation of the (**a**) AR- and (**b**) ER-mediated AOPs based on Tox21/ToxCast assays.

As shown in Figure 1a, the AR-mediated AOP included 20 *in vitro* assays, comprising three receptor binding assays (MIE), two co-regulator recruitment assays (KE1), one chromatin binding assay (KE2), two transcription factor regulation assays (KE3), five gene expression assays (KE4), two cell proliferation assays (KE5), and five cytotoxicity assays. The ER-mediated AOP (Figure 1b) encompassed 26 *in vitro* assays, including three receptor binding assays (MIE), six dimerization and co-regulator recruitment assays (KE1), two chromatin binding assays (KE2), four transcription factor activity assays (KE3), six assays for gene expression (KE4), one cell proliferation assay (KE5), and four cytotoxicity assays. Cytotoxicity assays were incorporated into both AOPs to provide critical context for interpreting agonistic and antagonistic responses, as they help discriminate true receptor-mediated activity from nonspecific effects driven by generalized cell stress.^35,36^

These assays were subsequently employed to train multitask learning models aimed at predicting AR- and ER-mediated responses. Modeling this dataset poses a substantial challenge, as the average imbalance ratio across training, validation, and test splits was 0.86, reflecting the dominance of inactives in the dataset (Figure 2a). To address this, four GNN architectures were explored (MPNN, GAT, GIN, and AttentiveFP), each optimized with virtual node embeddings, skip connections, and Jumping Knowledge mechanisms to improve chemical semantic representation and mitigate oversmoothing and oversquashing during training (Figure 2b). Performance evaluation revealed global AUROC values of 0.83 for MPNN, 0.81 for GAT, 0.78 for GIN, and 0.79 for AttentiveFP, with the multitask MPNN emerging as the most effective backbone for Tier-1 predictions (Figure 2c).

**Figure 2.**
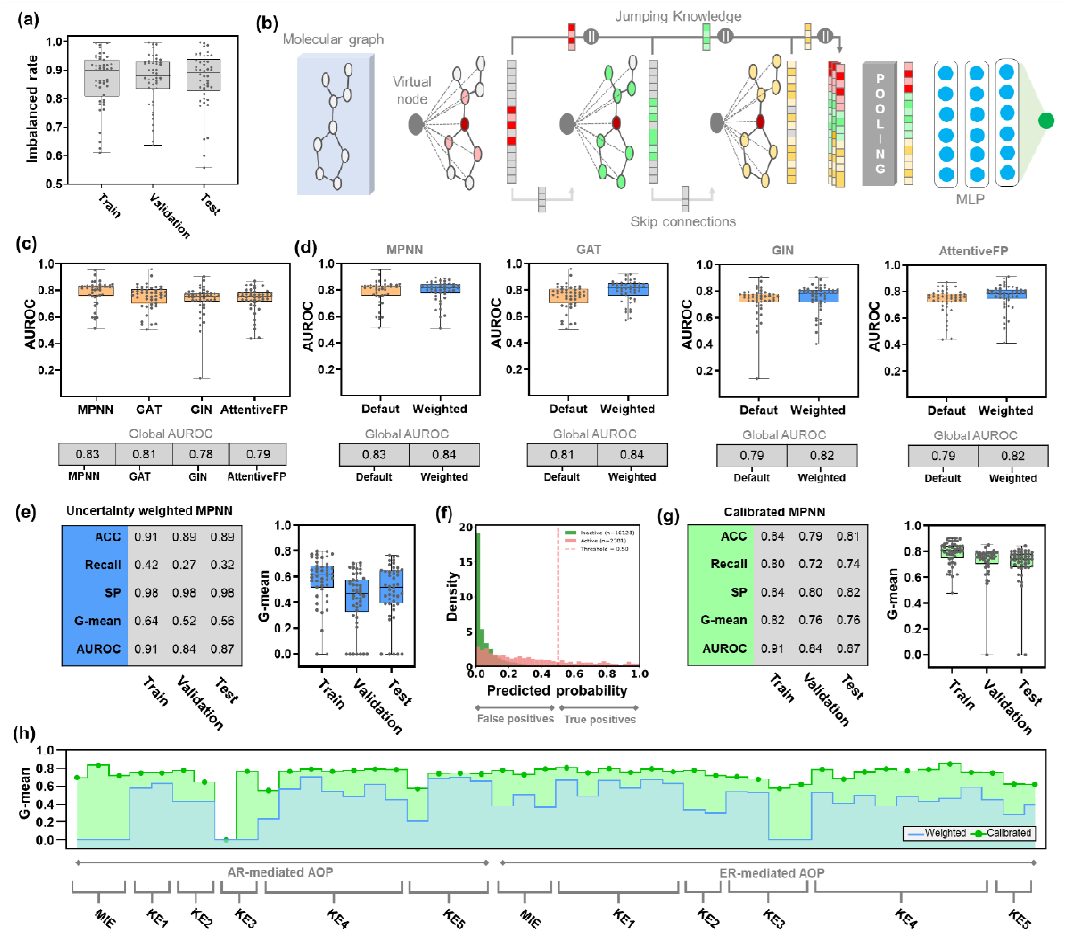
Statistical characteristics of multitask GNNs for predicting AR- and ER-mediated assay outcomes. (**a**) Imbalance ratios across training, validation, and test splits. (**b**) Schematic representation of multitask GNN architectures with virtual node, skip connections, and Jumping Knowledge mechanisms. (**c**) Global AUROC values of four GNN backbones. (**d**) Compariso of default vs. uncertainty-weighted loss across architectures. (**e**) Global performance metrics of the uncertainty-weighted MPNN and box plots of G-mean across tasks. (**f**) Predicted probability distributions showing class imbalance and suboptimality of the 0.5 threshold. (**g**) Global performance metrics of the calibrated MPNN with task-specific thresholds and box plots of G-mean across tasks. (**h**) Task-level G-mean before and after calibration across AR- and ER-mediated AOPs.

To further address heterogeneity across tasks, we implemented a homoscedastic uncertainty weighting framework to adaptively balance loss contributions during multitask learning. In this approach, each task is assigned a trainable variance parameter that downweights highly uncertain tasks, preventing them from dominating the optimization process. Meanwhile, a regularization term penalizes inflated variance estimates to avoid trivial solutions. As shown in Figure 2d, incorporating uncertainty weighting consistently improved performance across all architectures when compared with their default counterparts. The MPNN, which already achieved the best baseline performance (AUROC = 0.83), increased to 0.84 (+1.2%). The GAT exhibited the most significant relative gain, from 0.81 to 0.84 (+3.7%), while both GIN and AttentiveFP improved from 0.79 to 0.82 (+3.8%). These results indicate that homoscedastic uncertainty weighting provides a systematic advantage across diverse GNN backbones, with the strongest effect observed in models with lower baseline performance.

Figure 2e shows that the uncertainty-weighted MPNN maintained high accuracy across splits (ACC ≈0.89–0.91). However, data imbalance resulted in high specificity (SP = 0.98) but low recall (0.27–0.42), primarily reflecting the dominance of inactives. Consistently, G-mean values remained low (0.52–0.64) across splits, as highlighted in the accompanying box plots. The distribution of predicted probabilities (Figure 2f) further illustrates this bias, with inactives clustering near zero and most actives falling below the standard 0.5 decision boundary, highlighting that this default threshold is not optimal in such an imbalanced setting.

Probabilistic calibration^37^ was then applied to the best-performing model (MPNN) using task-specific thresholds optimized by G-mean. This calibration effectively shifts the decision boundaries on a task-by-task basis, moving them away from the default 0.5 threshold toward values that balance recall and SP. As a result, most active-class probability distributions undergo a leftward shift, recovering previously misclassified actives while preserving overall discrimination. As shown in Figure 2g, this strategy markedly increased recall (0.72–0.80) while maintaining balanced specificity (≈0.80–0.84). Consequently, G-mean values improved to 0.76–0.82 across splits, corresponding to relative gains of +28% (train), +46% (validation), and +36% (test), as illustrated in the box plots. At the individual assay level (Figure 2h), task-wise calibration yielded consistent improvements in G-mean across most endpoints within both AR- and ER-mediated AOPs. Only a few tasks showed limited or no gain, which is consistent with the high imbalance ratios and sparse representation of minority-class samples across the data splits in these assays.

### 2.2. Tier-2

Tier-2 modeling was designed to predict *in vivo* endocrine outcomes, specifically the Hershberger and uterotrophic assays, by integrating chemical features with pathway-derived predictions from Tier-1. The Hershberger dataset^38^ comprised 136 compounds, with activity defined by androgen-dependent changes in the weights of accessory sex tissues. The uterotrophic dataset^39^ comprised 118 compounds, where estrogenic activity was determined by measuring uterine weight changes in immature or ovariectomized female rodents. To expand the limited sample size of both assays, data augmentation was applied by incorporating experimentally confirmed inactives, consistently identified across all ToxCast/Tox21 assays (Hershberger: 123 compounds, uterotrophic: 136 compounds), as well as 13 non-toxicants from EDSP Tier-1 assessments. As shown in Figure 3a, the Hershberger dataset consisted of 69 positive and 203 negative samples, while the uterotrophic dataset comprised 64 positive and 203 negative samples. Notably, both datasets provided broad coverage of the chemical space represented in the *in vitro* ToxCast/Tox21 assays (Figure 3b), indicating that the compounds used for Tier-2 predictions are well distributed across the diversity of tested chemicals.

**Figure 3.**
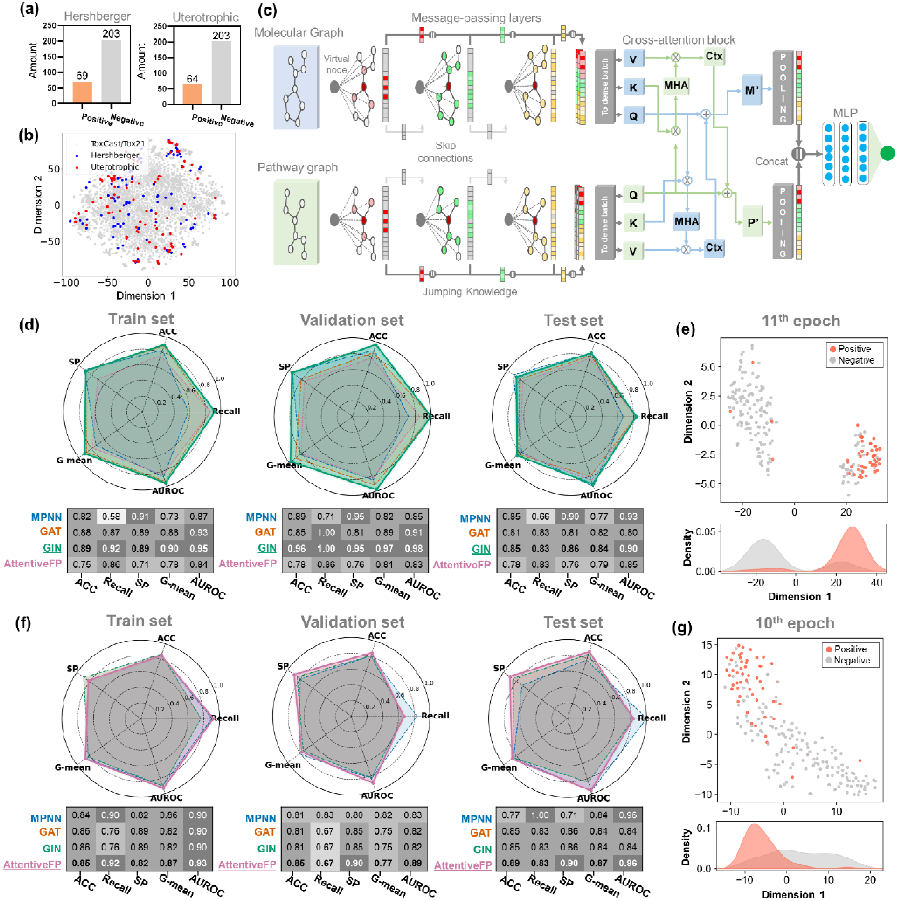
Statistical characteristics of multimodal GNN framework for predicting outcomes in Hershberger and uterotrophic assays. (**a**) Distribution of positive and negative samples in Hershberger and uterotrophic datasets after augmentation with consistently inactive compounds. (**b**) t-SNE projection of compounds showing broad coverage of the ToxCast/Tox21 chemical space. (**c**) Schematic representation of multimodal GNN architecture integrating molecular and pathway graphs through message-passing, cross-attention, and pooling. Predictive performance of multimodal GNN backbones (MPNN, GAT, GIN, AttentiveFP) on the (**d**) Hershberger and (**f**) uterotrophic assays across training, validation, and test sets. t-SNE embeddings of (**e**) Hershberger (GIN-based model, 11th epoch) and (**g**) uterotrophic predictions (AttentiveFP-based model, 10th epoch) showing clear separation of positive and negative compounds with corresponding KDE distributions.

To capture mechanistic context, pathway graphs were generated from AR- and ER-mediated AOPs, with nodes representing individual Tier-1 assays and edges reflecting their hierarchical position and co-occurrence patterns. These pathway graphs were integrated with molecular graphs through a multimodal GNN architecture (Figure 3c), where message-passing layers (AttentiveFP, GAT, GIN, and MPNN) processed each graph separately and bidirectional cross-attention blocks enabled information exchange between them. The refined molecular and pathway embeddings were then pooled, concatenated, and passed to a final classifier, providing joint representations optimized for predicting *in vivo* outcomes.

All multimodal GNN architectures demonstrated strong predictive performance for the Hershberger assay, with AUROC values consistently above 0.80 across validation and test (Figure 3d). Among them, the GIN-based model (probability threshold = 0.40) provided the best overall results, achieving AUROCs of 0.98 and 0.90, respectively. This model also reached the highest accuracy (0.96–0.85) and the best balance between Recall and SP, as reflected by G-mean values of 0.97–0.84. This superior performance was further supported by the t-SNE projection of learned embeddings (Figure 3e), which revealed a clear separation between positive (orange) and negative (gray) compounds. The corresponding KDE distributions confirmed minimal overlap between the two classes, highlighting the model’s ability to generate well-structured latent representations that discriminate against AR-mediated responses. Multimodal GNNs also showed consistently high predictive performance in the uterotrophic assay, achieving AUROC values greater than 0.82 across training, validation, and test sets (Figure 3f). The AttentiveFP-based model (probability threshold = 0.50) yielded the strongest results, reaching AUROCs of 0.89 and 0.96 for the validation and test sets, respectively. This model also maintained high accuracy (0.85–0.89) and delivered the most balanced trade-off between recall and specificity, as reflected by G-mean values of 0.77–0.87. The t-SNE projections of learned embeddings (Figure 3g) revealed distinct clustering of positive (orange) and negative (gray) compounds, with minimal overlap in KDE density, reinforcing the strength of AttentiveFP in modeling ER-mediated responses.

### 2.3. Ablation study

Ablation analysis was next performed to evaluate the contribution of each type of information (i.e., Tier-1 predictions, molecular graphs, and pathway graphs) to the predictive performance of Tier-2 models (Figure 4). Initially, the top-K algorithm was developed to assess whether Tier-1 predictions alone were sufficient for *in vivo* extrapolation. This approach classifies a compound as positive when the number of activated levels (MIEs and KEs) within the AOP exceeds a predefined threshold, reflecting the principle that progression across multiple key events increases the likelihood of an adverse outcome.^40,41^ As shown in Figure 4a, this approach yielded low performance, with an AUROC of 0.61 ± 0.14 for the Hershberger assay (−0.27 AUROC) and 0.63 ± 0.20 for the uterotrophic assay (−0.27 AUROC). These results suggest that *in vitro* outputs, when isolated, lack the resolution necessary to capture organism-level responses.

**Figure 4.**
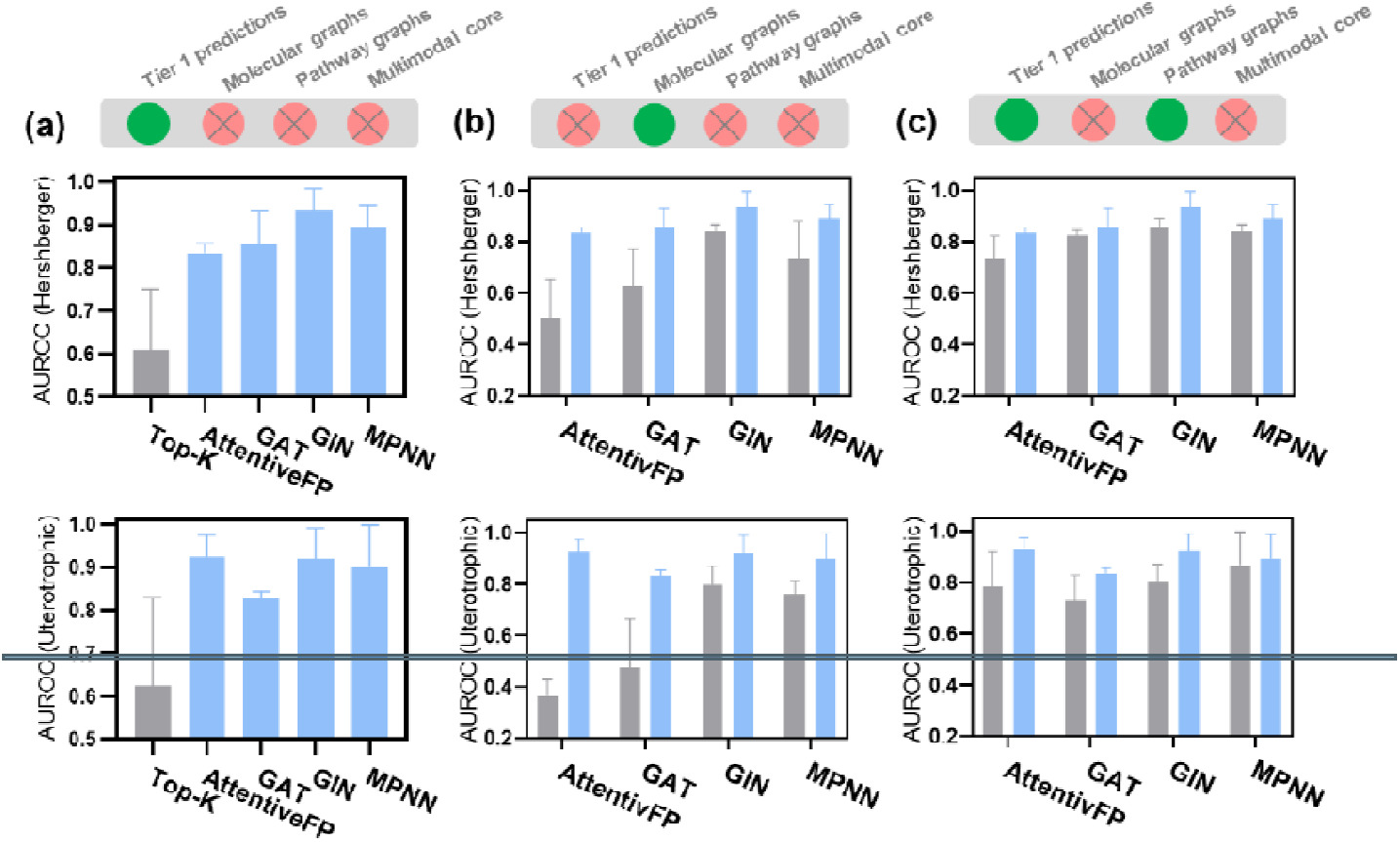
Ablation analysis of multimodal GNNs for predicting outcomes in Hershberger and uterotrophic assays. (**a**) AUROC evaluation of the top-K rule-based algorithm using Tier-1 predictions to classify compounds according to the number of activated AOP levels. (**b**) AUROC evaluation of the framework restricted to molecular graphs, isolating the contribution of chemical structure information. (**c**) AUROC evaluation of the framework restricted to pathway graphs, isolating the contribution of AOP-derived features. Error bars represent standard deviations across data splits.

#### Algorithm

Top-K algorithm for *in vivo* endocrine disruption classification

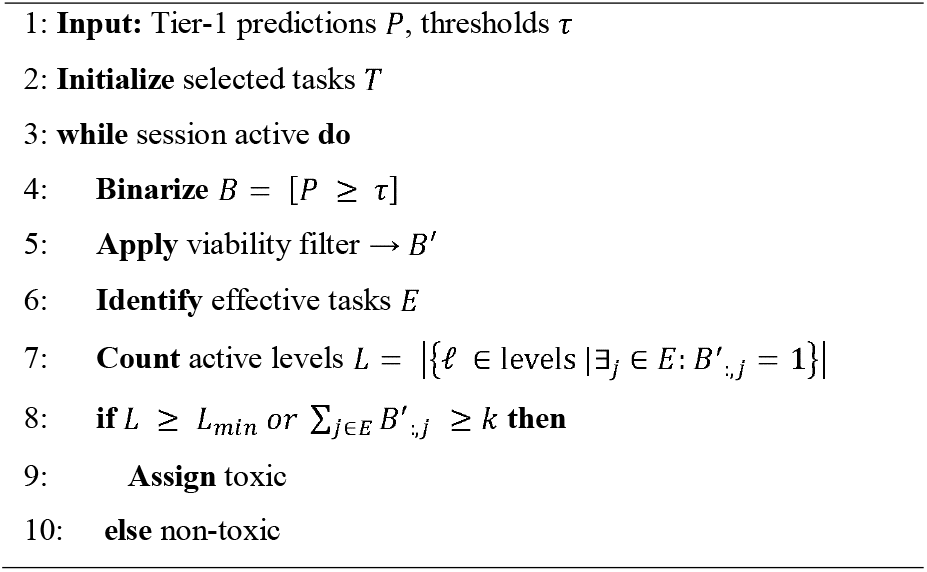

Subsequently, the contributions of molecular and pathway graphs were examined independently by deactivating each arm of the multimodal GNN framework. Models restricted to molecular graphs (Figure 4b) showed limited performance, with mean AUROC values of 0.68 ± 0.16 for the Hershberger assay (−0.20 AUROC) and 0.60 ± 0.21 for the uterotrophic assay (−0.30 AUROC), compared to the multimodal design (0.88 ± 0.06 and 0.90 ± 0.06, respectively). Pathway graph models (Figure 4c) yielded relevant prediction performances, achieving 0.82 ± 0.06 for the Hershberger assay (−0.06 AUROC) and 0.80 ± 0.10 for the uterotrophic assay (−0.10 AUROC); however, they underperformed relative to the multimodal framework. Overall, the combination of molecular and pathway graphs in the multimodal framework delivered the most accurate Tier-2 predictions.

### 2.4. External validation with literature compounds

To further interrogate the robustness and translational value of the Tier-2 models, an external validation was performed using compounds identified through a systematic literature review. Equivocal or conflicting reports, as well as mixtures and compounds associated with systemic effects such as changes in body weight, were excluded from the analysis. This evaluation included 19 chemicals with androgen-related outcomes directly relevant to the Hershberger assay and 26 chemicals with estrogen-related outcomes relevant to the uterotrophic assay (Tables 1 and 2). The selected compounds encompassed both substances tested in OECD guideline studies and others with well-documented *in vivo* outcomes strongly indicative of the same endocrine endpoints, thereby providing a stringent and diverse benchmark for model assessment.

**Table 1.**
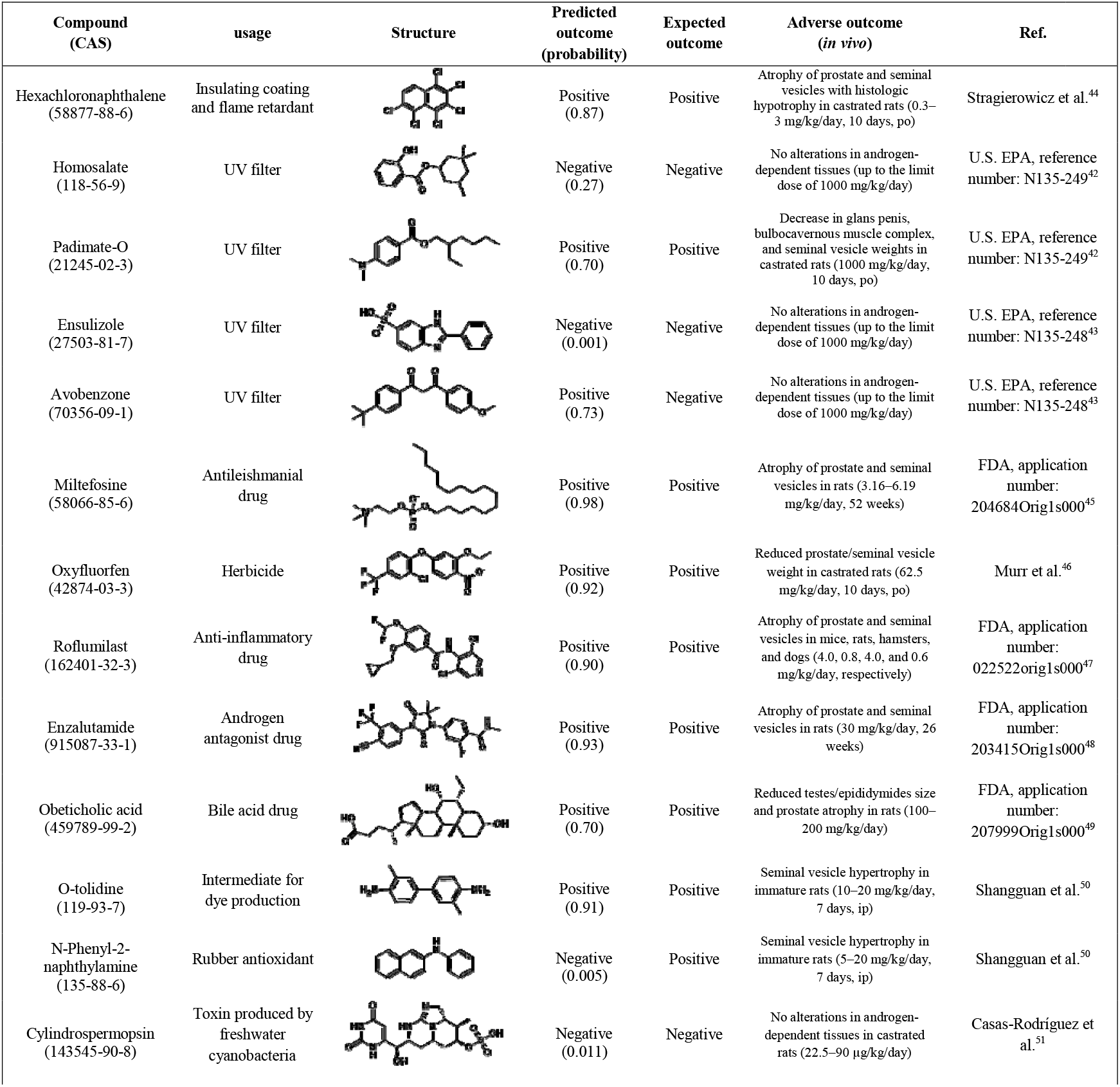
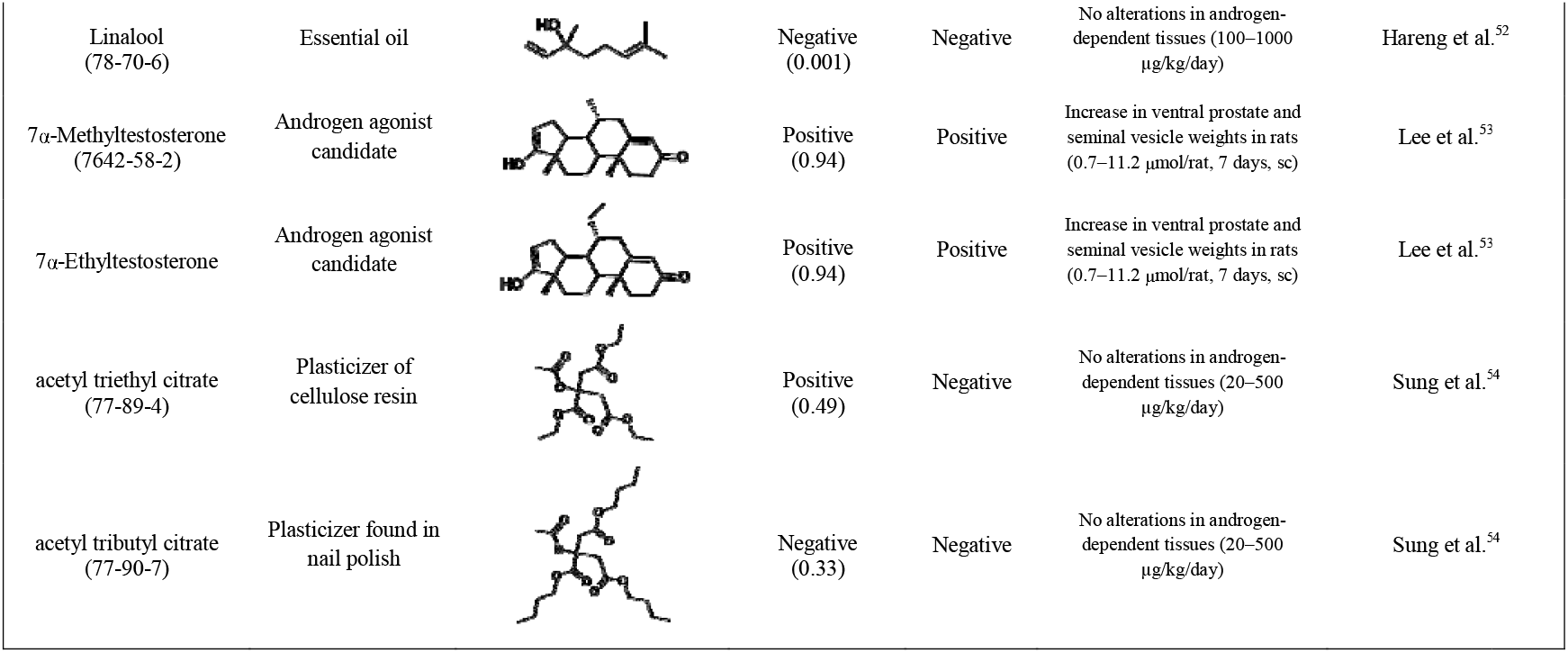
External validation of the Hershberger model with chemicals identified from the literature.

| Compound (CAS) | usage | Structure | Predicted outcome (probability) | Expected outcome | Adverse outcome (*in vivo*) | Ref. |
| --- | --- | --- | --- | --- | --- | --- |
| Hexachloronaphthalene (58877-88-6) | Insulating coating and flame retardant |  | Positive (0.87) | Positive | Atrophy of prostate and seminal vesicles with histologic hypotrophy in castrated rats (0.3–3 mg/kg/day, 10 days, po) | Stragierowicz et al.<sup>44</sup> |
| Homosalate (118-56-9) | UV filter |  | Negative (0.27) | Negative | No alterations in androgen-dependent tissues (up to the limit dose of 1000 mg/kg/day) | U.S. EPA, reference number: N135-249<sup>42</sup> |
| Padimate-O (21245-02-3) | UV filter |  | Positive (0.70) | Positive | Decrease in glans penis, bulbocavernous muscle complex, and seminal vesicle weights in castrated rats (1000 mg/kg/day, 10 days, po) | U.S. EPA, reference number: N135-249<sup>42</sup> |
| Ensulizole (27503-81-7) | UV filter |  | Negative (0.001) | Negative | No alterations in androgen-dependent tissues (up to the limit dose of 1000 mg/kg/day) | U.S. EPA, reference number: N135-248<sup>43</sup> |
| Avobenzone (70356-09-1) | UV filter |  | Positive (0.73) | Negative | No alterations in androgen-dependent tissues (up to the limit dose of 1000 mg/kg/day) | U.S. EPA, reference number: N135-248<sup>43</sup> |
| Miltefosine (58066-85-6) | Antileishmanial drug |  | Positive (0.98) | Positive | Atrophy of prostate and seminal vesicles in rats (3.16–6.19 mg/kg/day, 52 weeks) | FDA, application number: 204684Orig1s000<sup>45</sup> |
| Oxyfluorfen (42874-03-3) | Herbicide |  | Positive (0.92) | Positive | Reduced prostate/seminal vesicle weight in castrated rats (62.5 mg/kg/day, 10 days, po) | Murr et al.<sup>46</sup> |
| Roflumilast (162401-32-3) | Anti-inflammatory drug |  | Positive (0.90) | Positive | Atrophy of prostate and seminal vesicles in mice, rats, hamsters, and dogs (4.0, 0.8, 4.0, and 0.6 mg/kg/day, respectively) | FDA, application number: 022522Orig1s000<sup>47</sup> |
| Enzalutamide (915087-33-1) | Androgen antagonist drug |  | Positive (0.93) | Positive | Atrophy of prostate and seminal vesicles in rats (30 mg/kg/day, 26 weeks) | FDA, application number: 203415Orig1s000<sup>48</sup> |
| Obeticholic acid (459789-99-2) | Bile acid drug |  | Positive (0.70) | Positive | Reduced testes/epididymides size and prostate atrophy in rats (100–200 mg/kg/day) | FDA, application number: 207999Orig1s000<sup>49</sup> |
| O-tolidine (119-93-7) | Intermediate for dye production |  | Positive (0.91) | Positive | Seminal vesicle hypertrophy in immature rats (10–20 mg/kg/day, 7 days, ip) | Shangguan et al.<sup>50</sup> |
| N-Phenyl-2-naphthylamine (135-88-6) | Rubber antioxidant |  | Negative (0.005) | Positive | Seminal vesicle hypertrophy in immature rats (5–20 mg/kg/day, 7 days, ip) | Shangguan et al.<sup>50</sup> |
| Cylindrospermopsin (143545-90-8) | Toxin produced by freshwater cyanobacteria |  | Negative (0.011) | Negative | No alterations in androgen-dependent tissues in castrated rats (22.5–90 µg/kg/day) | Casas-Rodríguez et al.<sup>51</sup> |
| Linalool<br>(78-70-6) | Essential oil |  | Negative<br>(0.001) | Negative | No alterations in androgen-dependent tissues (100–1000 µg/kg/day) | Hareng et al.<sup>52</sup> |
| 7α-Methyltestosterone<br>(7642-58-2) | Androgen agonist candidate | | Positive<br>(0.94) | Positive | Increase in ventral prostate and seminal vesicle weights in rats (0.7–11.2 µmol/rat, 7 days, sc) | Lee et al.<sup>53</sup> |
| 7α-Ethyltestosterone | Androgen agonist candidate | | Positive<br>(0.94) | Positive | Increase in ventral prostate and seminal vesicle weights in rats (0.7–11.2 µmol/rat, 7 days, sc) | Lee et al.<sup>53</sup> |
| acetyl triethyl citrate<br>(77-89-4) | Plasticizer of cellulose resin |  | Positive<br>(0.49) | Negative | No alterations in androgen-dependent tissues (20–500 µg/kg/day) | Sung et al.<sup>54</sup> |
| acetyl tributyl citrate<br>(77-90-7) | Plasticizer found in nail polish |  | Negative<br>(0.33) | Negative | No alterations in androgen-dependent tissues (20–500 µg/kg/day) | Sung et al.<sup>54</sup> |

**Table 2.**
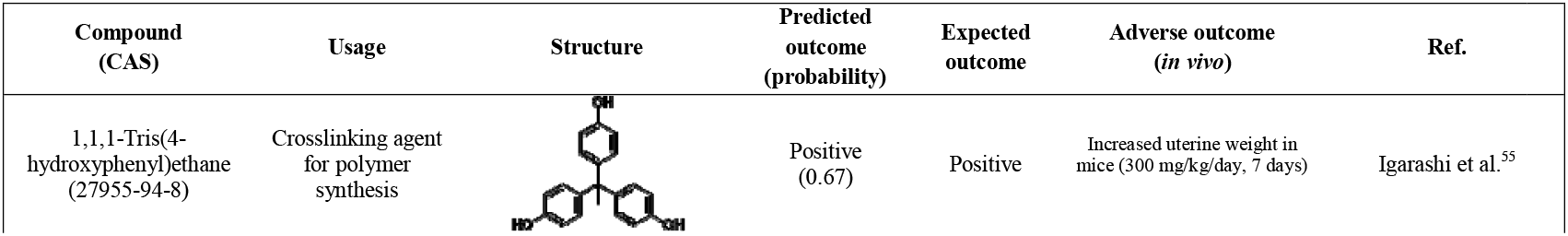
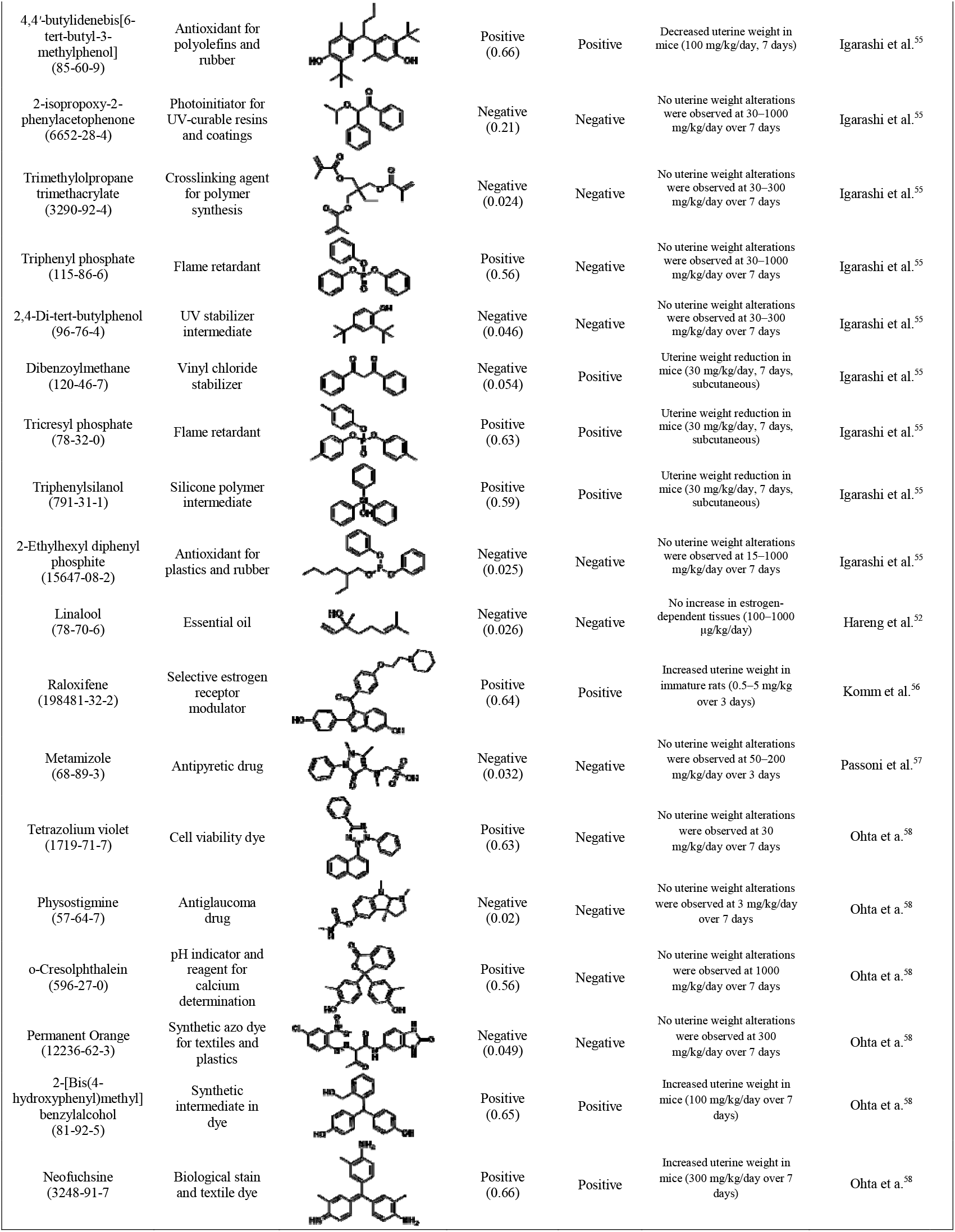
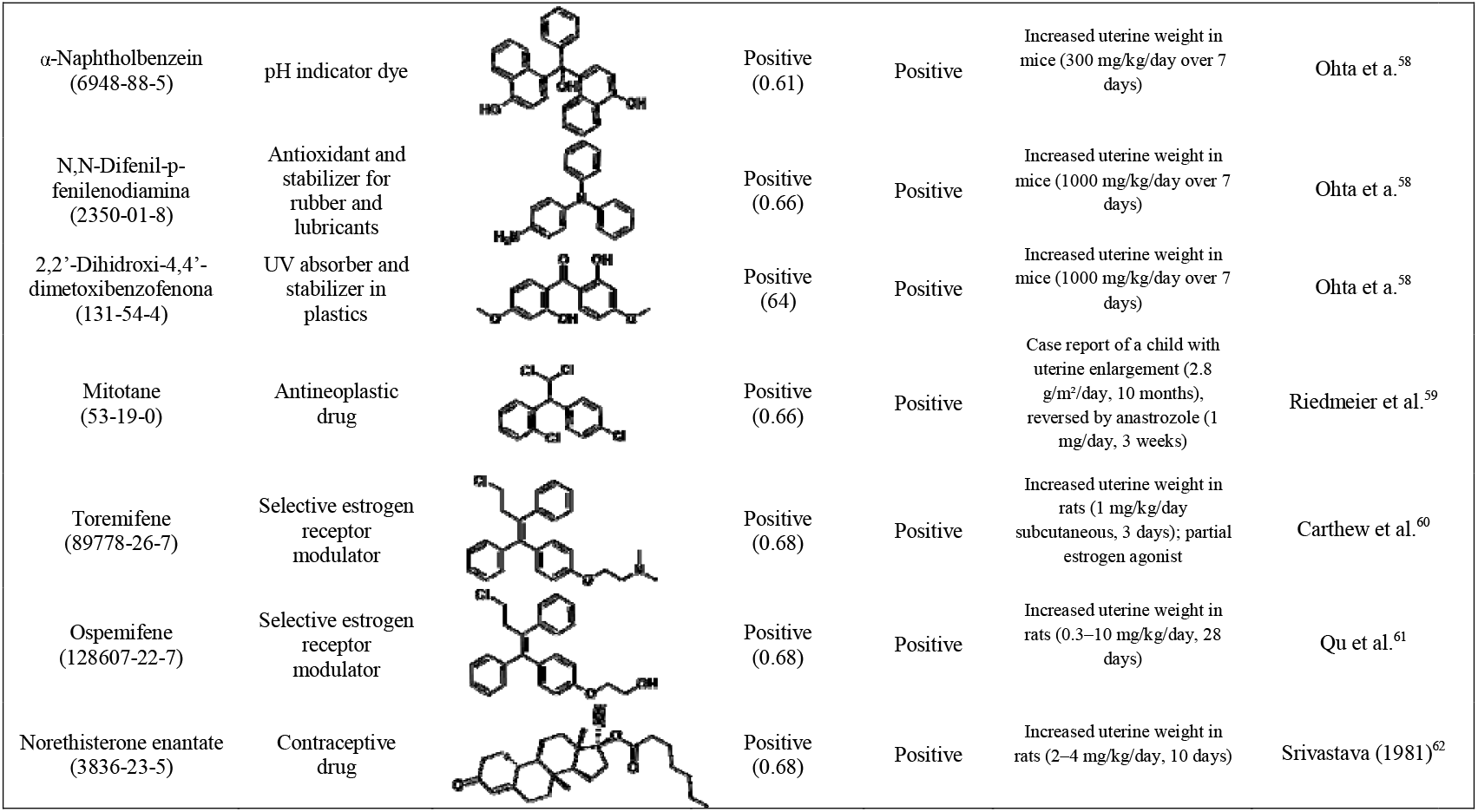
External validation of the uterotrophic model with chemicals identified from the literature.

| Compound<br>(CAS) | Usage | Structure | Predicted<br>outcome<br>(probability) | Expected<br>outcome | Adverse outcome<br>(*in vivo*) | Ref. |
| --- | --- | --- | --- | --- | --- | --- |
| 1,1,1-Tris(4-hydroxyphenyl)ethane<br>(27955-94-8) | Crosslinking agent<br>for polymer<br>synthesis |  | Positive<br>(0.67) | Positive | Increased uterine weight in mice (300 mg/kg/day, 7 days) | Igarashi et al.<sup>55</sup> |
| 4,4'-butylidenebis[6-tert-butyl-3-methylphenol]<br>(85-60-9) | Antioxidant for polyolefins and rubber |  | Positive<br>(0.66) | Positive | Decreased uterine weight in mice (100 mg/kg/day, 7 days) | Igarashi et al.<sup>55</sup> |
| 2-isopropoxy-2-phenylacetophenone<br>(6652-28-4) | Photoinitiator for UV-curable resins and coatings |  | Negative<br>(0.21) | Negative | No uterine weight alterations were observed at 30–1000 mg/kg/day over 7 days | Igarashi et al.<sup>55</sup> |
| Trimethylolpropane trimethacrylate<br>(3290-92-4) | Crosslinking agent for polymer synthesis |  | Negative<br>(0.024) | Negative | No uterine weight alterations were observed at 30–300 mg/kg/day over 7 days | Igarashi et al.<sup>55</sup> |
| Triphenyl phosphate<br>(115-86-6) | Flame retardant |  | Positive<br>(0.56) | Negative | No uterine weight alterations were observed at 30–1000 mg/kg/day over 7 days | Igarashi et al.<sup>55</sup> |
| 2,4-Di-tert-butylphenol<br>(96-76-4) | UV stabilizer intermediate |  | Negative<br>(0.046) | Negative | No uterine weight alterations were observed at 30–300 mg/kg/day over 7 days | Igarashi et al.<sup>55</sup> |
| Dibenzoylmethane<br>(120-46-7) | Vinyl chloride stabilizer |  | Negative<br>(0.054) | Positive | Uterine weight reduction in mice (30 mg/kg/day, 7 days, subcutaneous) | Igarashi et al.<sup>55</sup> |
| Tricresyl phosphate<br>(78-32-0) | Flame retardant |  | Positive<br>(0.63) | Positive | Uterine weight reduction in mice (30 mg/kg/day, 7 days, subcutaneous) | Igarashi et al.<sup>55</sup> |
| Triphenylsilanol<br>(791-31-1) | Silicone polymer intermediate |  | Positive<br>(0.59) | Positive | Uterine weight reduction in mice (30 mg/kg/day, 7 days, subcutaneous) | Igarashi et al.<sup>55</sup> |
| 2-Ethylhexyl diphenyl phosphite<br>(15647-08-2) | Antioxidant for plastics and rubber |  | Negative<br>(0.025) | Negative | No uterine weight alterations were observed at 15–1000 mg/kg/day over 7 days | Igarashi et al.<sup>55</sup> |
| Linalool<br>(78-70-6) | Essential oil |  | Negative<br>(0.026) | Negative | No increase in estrogen-dependent tissues (100–1000 µg/kg/day) | Hareng et al.<sup>52</sup> |
| Raloxifene<br>(198481-32-2) | Selective estrogen receptor modulator |  | Positive<br>(0.64) | Positive | Increased uterine weight in immature rats (0.5–5 mg/kg over 3 days) | Komm et al.<sup>56</sup> |
| Metamizole<br>(68-89-3) | Antipyretic drug |  | Negative<br>(0.032) | Negative | No uterine weight alterations were observed at 50–200 mg/kg/day over 3 days | Passoni et al.<sup>57</sup> |
| Tetrazolium violet<br>(1719-71-7) | Cell viability dye |  | Positive<br>(0.63) | Negative | No uterine weight alterations were observed at 30 mg/kg/day over 7 days | Ohta et al.<sup>58</sup> |
| Physostigmine<br>(57-64-7) | Antiglaucoma drug |  | Negative<br>(0.02) | Negative | No uterine weight alterations were observed at 3 mg/kg/day over 7 days | Ohta et al.<sup>58</sup> |
| o-Cresolphthalein<br>(596-27-0) | pH indicator and reagent for calcium determination |  | Positive<br>(0.56) | Negative | No uterine weight alterations were observed at 1000 mg/kg/day over 7 days | Ohta et al.<sup>58</sup> |
| Permanent Orange<br>(12236-62-3) | Synthetic azo dye for textiles and plastics |  | Negative<br>(0.049) | Negative | No uterine weight alterations were observed at 300 mg/kg/day over 7 days | Ohta et al.<sup>58</sup> |
| 2-[Bis(4-hydroxyphenyl)methyl]benzylalcohol<br>(81-92-5) | Synthetic intermediate in dye |  | Positive<br>(0.65) | Positive | Increased uterine weight in mice (100 mg/kg/day over 7 days) | Ohta et al.<sup>58</sup> |
| Neofuchsin<br>(3248-91-7) | Biological stain and textile dye |  | Positive<br>(0.66) | Positive | Increased uterine weight in mice (300 mg/kg/day over 7 days) | Ohta et al.<sup>58</sup> |
| α-Naphtholbenzein<br>(6948-88-5) | pH indicator dye | | Positive<br>(0.61) | Positive | Increased uterine weight in mice (300 mg/kg/day over 7 days) | Ohta et al.<sup>58</sup> |
| N,N-Difenil-p-fenilenodiamina<br>(2350-01-8) | Antioxidant and stabilizer for rubber and lubricants |  | Positive<br>(0.66) | Positive | Increased uterine weight in mice (1000 mg/kg/day over 7 days) | Ohta et al.<sup>58</sup> |
| 2,2'-Dihidroxi-4,4'-dimetoxibenzofenona<br>(131-54-4) | UV absorber and stabilizer in plastics |  | Positive<br>(.64) | Positive | Increased uterine weight in mice (1000 mg/kg/day over 7 days) | Ohta et al.<sup>58</sup> |
| Mitotane<br>(53-19-0) | Antineoplastic drug |  | Positive<br>(0.66) | Positive | Case report of a child with uterine enlargement (2.8 g/m<sup>2</sup>/day, 10 months), reversed by anastrozole (1 mg/day, 3 weeks) | Riedmeier et al.<sup>59</sup> |
| Toremifene<br>(89778-26-7) | Selective estrogen receptor modulator |  | Positive<br>(0.68) | Positive | Increased uterine weight in rats (1 mg/kg/day subcutaneous, 3 days); partial estrogen agonist | Carthew et al.<sup>60</sup> |
| Ospemifene<br>(128607-22-7) | Selective estrogen receptor modulator |  | Positive<br>(0.68) | Positive | Increased uterine weight in rats (0.3–10 mg/kg/day, 28 days) | Qu et al.<sup>61</sup> |
| Norethisterone enantate<br>(3836-23-5) | Contraceptive drug |  | Positive<br>(0.68) | Positive | Increased uterine weight in rats (2–4 mg/kg/day, 10 days) | Srivastava (1981)<sup>62</sup> |

Application of the Tier-2 framework to this independent dataset revealed a high degree of concordance between predicted and observed outcomes. The Hershberger model correctly classified 15 of 18 compounds (ACC = 83%), while the uterotrophic model accurately predicted 22 of 26 cases (ACC = 84%). Among the compounds evaluated, specific examples illustrate the predictive capacity of the Tier-2 framework. Padimate-O (CAS no. 21245-02-3), a UV filter, was correctly identified as positive by the model (probability = 0.70). Consistent with experimental evidence, *in vivo* studies report decreases in glans penis, bulbocavernous muscle complex, and seminal vesicle weights in rats exposed orally at 1000 mg/kg/day for 10 days.^42^ In contrast, ensulizole (CAS no. 27503-81-7), another UV filter, was correctly predicted as negative (probability = 0.001), in agreement with Hershberger assays, which showed no alterations in androgen-dependent tissues up to the limit dose of 1000 mg/kg/day.^43^

The application of the Tier-2 framework to the uterotrophic assay further illustrates the alignment between predicted outcomes and experimental evidence. The compound 4,4′-Butylidenebis[6-tert-butyl-3-methylphenol] (CAS no. 85-60-9), an antioxidant widely used in polyolefins and rubber applications, was correctly predicted as positive for uterotrophic activity (probability = 0.66). Experimental evidence supported this prediction, with *in vivo* uterotrophic studies reporting a significant decrease in uterine weight in mice following exposure at 100 mg/kg/day for 7 days.^55^ In contrast, 2-Ethylhexyl diphenyl phosphite (CAS no. 15647-08-2), an antioxidant commonly employed in plastics and rubber manufacturing, was predicted as negative for uterotrophic activity (probability = 0.025). This prediction was consistent with *in vivo* data, as no alterations in uterine weight were detected in mice exposed to doses ranging from 15 to 1000 mg/kg/day over a 7-day period.^55^ These results demonstrate that the multimodal cross-attentive GNNs extend beyond internal validation, effectively generalizing to novel compounds with organism-level endocrine effects. Such consistency with independent *in vivo* evidence underscores the reliability of the framework and highlights its potential to inform regulatory evaluation where traditional testing remains limited.

### 2.5. Tier-2 explainability

Besides assessing the model’s performance, it is often beneficial to examine the “black box” of the trained model to gain a deeper understanding of which molecular substructures and pathway-level assays most strongly drive *in vivo* endocrine disruption. To enable explainability, the present framework incorporates two original strategies designed in this study: cross-attention mapping and counterfactual perturbations. Cross-attention analysis was performed bidirectionally, mapping molecular substructures to pathway assays (M2P) and, conversely, linking pathway nodes back to their most influential molecular features (P2M). This dual perspective enabled the identification of explicit connections between atomic features and AR- and ER-mediated assay nodes, highlighting how chemical motifs align with pathway-level perturbations. Complementing this mechanistic mapping, counterfactual perturbations were employed to generate alternative scenarios, enabling the evaluation of how molecular modifications or pathway disruptions may alter downstream outcomes. To illustrate how the Tier-2 framework provides mechanistic interpretability for endocrine disruption predictions, we first examined two representative compounds (Figure 5): hexachloronaphthalene (CAS no. 58877-88-6), classified as positive in the Hershberger assay, and 1,1,1-tris(4-hydroxyphenyl)ethane (CAS no. 27955-94-8), identified as positive in the uterotrophic assay.

**Figure 5.**
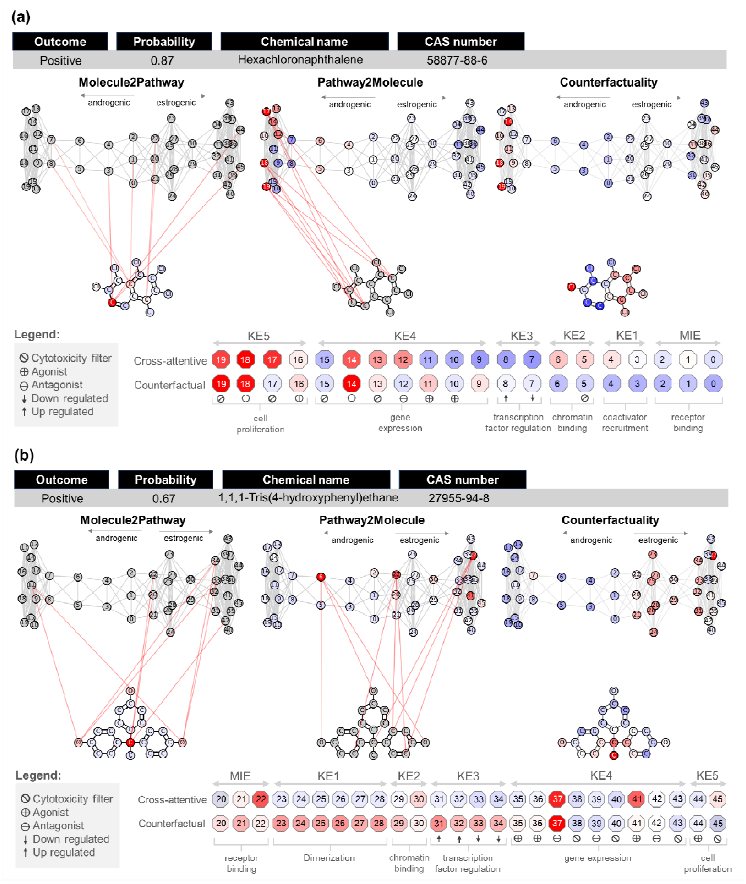
Integrative cross-attention (non-causal) and counterfactual analyses illustrating the mechanistic interpretability of Tier-2 predictions for (**a**) hexachloronaphthalene (Hershberger positive) and (**b**) 1,1,1-tris(4-hydroxyphenyl)ethane (uterotrophic positive).

The model’s classification of hexachloronaphthalene as a positive compound in the Hershberger assay is attributed to its function as an AR antagonist (probability = 0.87). The primary evidence for this was established through a cross-attention analysis, which elucidated the mechanistic underpinnings of the prediction (Figure 5a). This analysis identified strong, bidirectional connections between the molecule’s chlorinated aromatic motifs and specific nodes within the AR-mediated pathway. Such a connection confirms a robust structure-activity relationship, indicating that these structural features are integral to the observed toxicological activity. This conclusion was further substantiated by counterfactual perturbations, which demonstrated that computational modification of the chlorinated ring system would negate the predicted outcome, thereby confirming the structural feature’s causal role in the compound’s antagonistic profile.

The model’s reasoning is further clarified by tracing the compound’s influence along the AOP. The analysis assigned a negative influence (indicated by a blue color) to early-stage events, including the MIE of receptor binding and subsequent key events, such as coactivator recruitment (KE1) and transcription factor regulation (KE3). This pattern suggests the compound does not operate via a classical agonistic mechanism that would activate the pathway from its inception. In contrast, the model identified a strong positive influence (indicated by a red color) concentrated in downstream, antagonistic assays. These correspond to KE4, which involves the regulation of androgen-dependent gene expression, and KE5, which pertains to cell proliferation. The model thus inferred an antagonistic mechanism from this distinct AOP signature, characterized by the absence of initial pathway activation coupled with a significant blockage of terminal biological functions.

The mechanistic insights derived from the model show a high degree of concordance with experimental *in vivo* data. Scientific literature reports that exposure to hexachloronaphthalene results in prostate and seminal vesicle atrophy in rats, a hallmark of AR antagonism. The model’s identification of pathway disruption at KE4 and KE5 provides the direct mechanistic explanation for this adverse outcome. Androgen-dependent tissues require continuous signaling through the AR to maintain cellular homeostasis and mass. By blocking this pathway, hexachloronaphthalene inhibits the transcription of essential androgen-dependent genes (disruption of KE4). The subsequent failure of cells to proliferate and survive (disruption of KE5) leads to a net loss of tissue mass, which is physically observed as atrophy. This causal chain, validated by the findings of Stragierowicz et al.,^44^ directly links the model’s AOP-based prediction to the empirically observed *in vivo* endpoint, supporting the compound’s classification as positive in the Hershberger model.

The model classifies 1,1,1-Tris(4-hydroxyphenyl)ethane as a positive compound for uterotrophic activity, providing a mechanistic rationale for its 0.67 prediction probability. Cross-attention analysis identified the molecule’s aromatic hydroxyl (-OH) groups as the primary structural determinants of its bioactivity. Strong, bidirectional connections were established between these functional moieties and nodes within the estrogen receptor (ER)-mediated pathway (Figure 5b). This indicates that the model not only found a correlation but also identified these hydroxyl groups as central to the compound’s mechanism of action, directly linking chemical structure to predicted toxicological function.

An examination of the model’s reasoning along the AOP reveals a nuanced and complex mechanistic signature. The prediction was not driven by a simple agonistic signal; instead, the model weighed conflicting evidence. A positive influence (red) was attributed to antagonistic assays at the MIE, KE2 (chromatin binding), and a mixed agonist/antagonist profile at KE4 (gene expression). This was contrasted with a negative influence (blue) at KE1 (dimerization) and KE5 (cell proliferation). This complex pattern aligns with the “dual behavior” of certain estrogenic compounds, which can exhibit both agonistic and antagonistic features depending on the toxicological context, similar to Selective Estrogen Receptor Modulators (SERMs).^63^ Counterfactual analysis further layered this complexity, showing that KE3 (transcription-factor regulation) and other KE4 assays also contributed significantly to the positive prediction.

A compelling validation of the model’s assessment is found in its alignment with the definitive *in vivo* endpoint. Experimental evidence from Igarashi et al.^55^ confirms that the administration of 1,1,1-Tris(4-hydroxyphenyl)ethane induces a significant increase in uterine weight in ovariectomized mice, the classical uterotrophic response. This demonstrates that, despite the partial antagonistic signals detected across the AOP, the compound’s net physiological effect in uterine tissue is predominantly agonistic, resulting in cellular proliferation and tissue growth. The model’s ability to process these seemingly contradictory signals and still forecast the correct organism-level outcome showcases its capacity to learn complex, non-linear biological signatures and accurately integrate them into a final, validated prediction.

## 3. CONCLUSIONS

This work introduces an innovative multimodal cross-attentive framework that fuses chemical graph information with AOP-anchored, assay-level signals to predict organism-level endocrine outcomes. The multi-tier design, composed of multitask Tier-1 models that learn AR/ER MIEs and KEs, followed by Tier-2 cross-attentive models, achieved high and consistent predictive performance for both Hershberger and uterotrophic endpoints. It outperformed single-source baselines in ablation studies and generalized well in external validations. To render predictions interpretable, cross-attention analysis is performed bidirectionally, mapping molecular substructures to pathway assays (M2P) and, conversely, linking pathway nodes back to their most influential molecular features (P2M). This dual perspective establishes explicit correspondences between atomic features and AR- and ER-mediated assay nodes across the AOP (MIE→KE5), clarifying how chemical motifs align with pathway-level perturbations. Coupled with counterfactual perturbations that quantify the impact of targeted molecular edits or pathway shocks on the predicted outcome, the framework identifies the assays that most determine each *in vivo* decision and the structural elements that drive those signals. In summary, this computational approach represents an early IATA-compatible tool for screening and prioritization, helping identify substances with the highest expected decision impact for further experimental evaluation. Its use can substantially reduce redundant animal testing and optimize resource allocation, while still acknowledging key limitations, such as the chemical space covered, the lack of metabolic competence in many *in vitro* sources, potential species- and strain-specific differences, untested dose–response regimens, and the inability to fully account for mixture effects. By making such constraints explicit, the framework facilitates evidence-based integration with regulatory testing strategies, thereby contributing to a more efficient and mechanistically informed risk assessment process.

## 4. METHODS

### 4.1. Data collection and curation

A dataset was compiled from the U.S. EPA ToxCast/Tox21 program (invitroDB v4.1), encompassing 46 *in vitro* assays relevant to AR- and ER-mediated AOPs. For the AR pathway, 18 assays were included, covering receptor binding, co-regulator recruitment, transcription factor activity, gene expression, and cell proliferation. For the ER pathway, 22 assays were considered, including receptor binding, dimerization and co-regulator recruitment, chromatin binding, transcription factor activity, gene expression, and cell proliferation. Cytotoxicity assays were also incorporated to interpret antagonistic activity. In parallel, *in vivo* data were obtained from rodent uterotrophic^39^ and Hershberger^38^ assays. To mitigate data scarcity, datasets were augmented with additional inactive chemicals from ToxCast/Tox21 and the U.S. EPA EDSP Tier-1 assessments. Subsequently, chemical structures in SMILES format and their bioactivity records were curated according to established best practices.^64,65^ The process involved standardization of functional groups and aromatic systems and removal of salts, mixtures, polymers, and organometallics. Duplicate entries with conflicting activity outcomes were excluded, while concordant duplicates were merged into a single representative record. Subsequently, the datasets were randomly split into training, validation, and test sets.

### 4.2. Tier-1 models

Tier-1 models were developed to predict the outcomes of 46 *in vitro* assays encompassing AR- and ER-mediated MIEs and KEs, using molecular graphs as feature representations.

#### 4.2.1. Molecular graphs

Molecular graphs were constructed from SMILES strings using RDKit v.2024.4.5, representing atoms as nodes and bonds as edges. Atom-level descriptors comprised one-hot encodings of atomic number, degree, formal charge, hybridization, and aromaticity. Bond features were encoded through one-hot representations of bond type, conjugation, ring membership, stereochemistry, and chirality.

#### 4.2.2. Architecture

The MPNN, GAT, and GIN architectures were refined through systematic incorporation of skip connections, virtual nodes, and Jumping Knowledge (JK) mechanisms to enhance chemical semantics. Furthermore, AttentiveFP architecture was employed without modification, as it inherently incorporates multiscale mechanisms analogous to those introduced in the optimized models. Skip connections were incorporated to mitigate over-smoothing and gradient degradation during deep message-passing as follows:

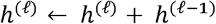

where *h*^(*ℓ* − 1)^, *h*^(*ℓ*)^ ∈ ℝ^d^are input and output features at layer *ℓ*. This additive transform acts as an identity initialization, preserving the gradient flow.

The virtual node mechanism was incorporated into each molecular graph to enrich chemical semantics by forging long-range connections among pharmacophoric and structural features. This node is initialized with a learnable embedding by 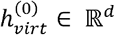 and interacts with all nodes by contributing additively to their representations at each message-passing layer. For a given graph *G* = (*V, E*), the representation of each atom *v* ∈ *V* at layer ℓ is modified as follows:

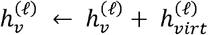

where 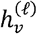 is the atom’s representation at layer ℓ, and 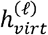 is the virtual node’s representation at the same layer. After message-passing, the virtual node embedding is updated by aggregating the current node states across graph:

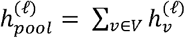 where 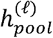 is the graph-level summary at layer ℓ, computed as permutation-invariant sum over all atomic representations 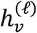. The pooled representation is then transformed through a learnable nonlinear function *MLP*^(*ℓ*)^, typically consisting of two linear layers with intermediate activation and normalization, to produce a virtual node embedding as follows:

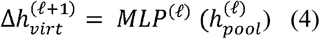

Finally, the virtual node embedding is updated by residual addition:

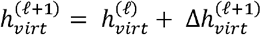

where the virtual node embedding from the previous layer 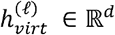 is incremented by the newly computed update 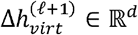.

The JK mechanism was incorporated into architectures to address this limitation by allowing the model to adaptively aggregate information from multiple message-passing depths, effectively capturing signals at varying neighborhood ranges. Formally, let H^(0),^ H^(1)^, …, H^(*L*)^ denote the sequence of node embeddings obtained at each of the *L* message-passing layers. JK aggregation builds a unified node representation through feature concatenation:

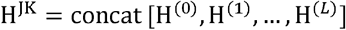

#### 4.2.3. Homoscedastic uncertainty

A homoscedastic uncertainty weighting framework,^66^ was employed to balance task contributions during multitask learning by accounting for task-dependent variance. For each task *t*, a trainable parameter 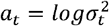 defined as the logarithm of the task-dependent variance was introduced to adaptively scale the classification loss as follows:

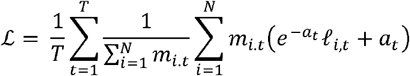

where 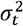 represents the homoscedastic uncertainty associated with task *t, T* is the number of tasks and *N* the number of samples, and individual task loss *ℓ*_*i,t*_ corresponds to the binary cross-entropy. The binary mask *m*_*i.t*_ ∈ {0,1} indicates whether a valid label is available for sample *i* in task *t*, ensuring that only valid entries contribute to the loss. The exponential term 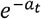 adaptively downweights tasks with higher uncertainty, while the additive *a*_*t*_ term penalizes overly large uncertainty estimates, preventing trivial solutions.

#### 4.2.4. Probabilistic calibration

Probabilistic calibration was applied in class-imbalanced settings using a threshold-moving strategy^37^ to adjust the decision boundaries based on raw predicted probabilities. Thresholds were evaluated individually for each MTL task across a 0.0–1.0 range with 0.01 increments on the training and validation sets, and optimal values were defined by the highest G-mean as follows:

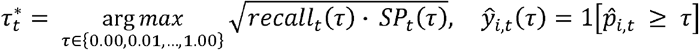

where *recall*_*t*_(*τ*) and *S*_*t*_(*τ*) denote recall and specificity, computed at threshold *τ* for task *t*. The indicator function 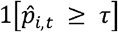 assigns a binary label *ŷ*_*i,t*_ (*τ*)to each prediction 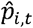, with outcome 1 if the predicted probability is greater than or equal to τ, and 0 otherwise. The calibrated thresholds 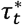 were subsequently applied to the test set to reclassify predictions, enhancing the trade-off between Recall and SP while maintaining AUROC.

### 4.3. Tier-2 models

Tier-2 models were developed to predict adverse outcomes from the *in vivo* Hershberger and uterotrophic assays, integrating molecular and pathway graphs as complementary feature representations.

#### 4.3.1. Pathway graphs

Pathway graphs were constructed from AR- and ER-mediated AOPs using hierarchical and co-occurrence information derived from the ToxCast/Tox21 dataset. Assays were grouped according to their MIE and KE classification, with rule levels assigned to reflect their position within AOPs. Nodes represented individual assays, annotated with Tier-1 logits, which are the raw outputs of the predictive model that quantify the strength and direction of the prediction. Edges were defined based on both structural hierarchy (linking events within the same pathway group and adjacent MIE/KE levels) and statistical associations derived from binarized Tier-1 model predictions. Association weights were computed using normalized pointwise mutual information (nPMI) and Yule’s Q values, calculated from predicted class co-occurrence patterns, and incorporated as edge attributes. The nPMI between tasks *a* and *b* was given by:

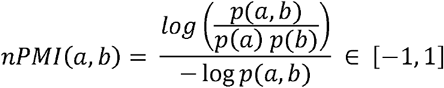

where *p* (*a, b*) is the joint probability of both tasks being predicted as positive, and *p* (*a*) and *p* (*b*) are their respective marginal probabilities. In parallel, Yules’s Q values derived from the odds ratio was computed as follows:

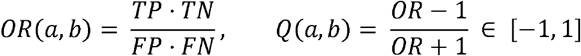

where TP, TN, FP, and FN denote the counts of true positives, true negatives, false positives and false negatives for the pair (*a, b*).

#### 4.3.2. Architecture

Four multimodal GNNs were developed using MPNN, GAT, GIN, and AttentiveFP message-passing mechanisms. Each model employed parallel message-passing arms for the molecular and pathway graphs, enhanced with skip connections, virtual nodes, and JK mechanisms. The outputs of both encoders were projected into a shared latent dimension and integrated through a bidirectional cross-attention mechanism. Multi-head attention (MHA) was computed as:

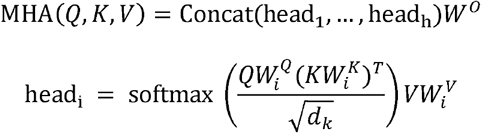

where Q, K, and V represent query, key, and value projections of the input node, *d*_*k*_ is the key dimension, and 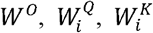, and 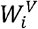 are learned projection matrices. In the M2P direction, pathway embeddings were updated as:

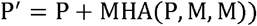

where P denotes the input pathway node embeddings and M the molecular embeddings. Conversely, in the P2M direction, molecular representations were updated as:

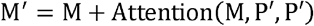

where P′ are the updated pathway embeddings. Finally, the refined embeddings P′ and M′ were pooled, concatenated, and passed through a multilayer perceptron to generate *in vivo* predictions

### 4.4. Model training and evaluation

All models were implemented in PyTorch v.2.5.1 with PyTorch Geometric v.2.6.1 and trained on an NVIDIA TITAN Xp GPU. Training was capped at 2,000 epochs using binary cross-entropy (BCE) loss, with early stopping applied when no improvement in validation loss was observed within a five-epoch adaptive window. A ReduceLROnPlateau scheduler was employed to reduce the learning rate when validation loss stagnated beyond the defined patience. To enhance efficiency, pruning techniques were integrated to halt underperforming trials. Hyperparameter tuning was performed using Optuna v.4.2.0, with 200 trials per architecture to systematically explore the parameter space. Model performance was assessed using ACC, recall, SP, and AUROC, with G-mean additionally considered to address class imbalance. Embedding vectors generated during training were retained and later projected via t-SNE and KDE to evaluate the structuring of the latent space across epochs.

### 4.5. Cross-attentive explainability

Cross-attentive explanations were generated by extracting bidirectional attention matrices between the molecular and pathway graphs during inference, where M2P and P2M attention weights were used to compute node-level importance scores. Formally, the importance score for a source node *i* was calculated as:

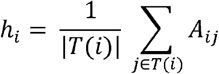

where *A*_*ij*_ denotes the cross-attention weight from node *i* to target *j*, and *T*(*i*) is the set of valid attended targets. In the M2P direction, *i* corresponds to a molecular node (atom) and *j* to a pathway node (assay). Conversely, in the P2M direction, *j* denotes a pathway node and *j* a molecular node. Interpretability was enhanced by retaining only the top-*k* attention links between *j* and *j*. Finally, both graphs were jointly rendered within the same spatial frame, with bidirectional cross-attention links superimposed to highlight the most influential molecular–assay interactions.

### 4.6. Counterfactual explainability

Stochastic perturbations were applied to the molecular and pathway graphs using rule-based isosteric and valence-preserving substitutions, and quasi-random assay perturbations derived from low-discrepancy sequences, respectively. Subsequently, the importance score of each node *i* was calculated as follows:

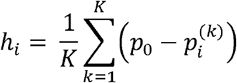

where *P*_0_ is the predicted probability for the unperturbed graph, 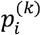 is the predicted probability after *k*-th perturbation of node *i*, and *K* is the number of perturbation samples. Graph-based clustering was then performed using connected components to group structurally connected atoms with similar contribution signals, with intra-cluster enhancement applied as:

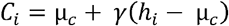

where *C*_*i*_ is the cluster-enhanced contribution score for atom 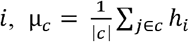 is the mean contribution *c*, and γ > 0 is the cluster-enhancement factor. Laplacian smoothing was subsequently applied within each cluster to reduce noise while preserving local structure as follows:

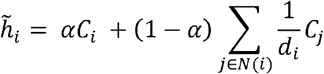

where *C*_*j*_ is the cluster-enhanced score of each neighbor *j* ∈ *N*(*i*), whereas *α* ∈ [0, 1] balances self-vs-neighbor contributions, *N*(*i*) is the set of neighbors of atom *i*, and *d*_*i*_= |*N*(*i*)| its degree. Finally, the 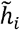 values are normalized (zero mean, unit variance) and projected onto 2D graph coordinates for visualization.

## Acknowledgements

The authors thank CNPq, CAPES, and FAPEG for financial support and fellowships. This study received financial support from the FAPEG (grant # 202310267001412), CAPES (Finance Code 001), and INCT (grant #408678/2024-0). BJN is CNPq productivity fellow (grant #311100/2023-6).

